# Different SUMO Paralogs Determine the Fate of WT and Mutant CFTRs: Biogenesis vs. Degradation

**DOI:** 10.1101/340851

**Authors:** Xiaoyan Gong, Yong Liao, Annette Ahner, Mads Breum Larsen, Xiaohui Wang, Carol A. Bertrand, Raymond A. Frizzell

**Affiliations:** Departments of Pediatrics and Cell Biology, University of Pittsburgh School of Medicine, Rangos Research Center, 4401 Penn Avenue, Pittsburgh, PA 15224

## Abstract

A pathway for CFTR degradation is initiated by Hsp27 which cooperates with Ubc9 and binds to the common F508del mutant to modify it with SUMO-2/3. These SUMO paralogs form poly-chains, which are recognized by the ubiquitin ligase, RNF4, for proteosomal degradation. Here, protein array analysis identified the SUMO E3, PIAS4, which increased WT and F508del CFTR biogenesis in CFBE airway cells. PIAS4 increased immature CFTR three-fold and doubled expression of mature CFTR, detected by biochemical and functional assays. In cycloheximide chase assays, PIAS4 slowed immature F508del degradation 3-fold and stabilized mature WT CFTR at the PM. PIAS4 knockdown reduced WT and F508del CFTR expression by 40-50%, suggesting a physiological role in CFTR biogenesis. PIAS4 modified F508del CFTR with SUMO-1 *in vivo* and reduced its conjugation to SUMO-2/3. These SUMO paralog specific effects of PIAS4 were reproduced *in vitro* using purified F508del NBD1 and SUMOylation reaction components. PIAS4 reduced endogenous ubiquitin conjugation to F508del CFTR by ~50%, and blocked the impact of RNF4 on mutant CFTR disposal. These findings indicate that different SUMO paralogs determine the fates of WT and mutant CFTRs, and they suggest that a paralog switch during biogenesis can direct these proteins to different outcomes: biogenesis vs. degradation.

## INTRODUCTION

The cystic fibrosis transmembrane conductance regulator (CFTR) is the basis of the cAMP/PKA-stimulated anion conductance at the apical membranes of secretory epithelial cells in the airways, intestines, pancreas and other systems (Frizzell and Hanrahan, 2012). As a member of the ABC transporter family, CFTR is composed of two membrane spanning domains (MSD1 and MSD2), two nucleotide binding domains (NBD1 and NBD2) and a unique and unstructured regulatory (R) domain. The R domain contains sites whose kinase-mediated phosphorylation enables CFTR channel gating via ATP binding and hydrolysis at the NBDs. The omission of phenylalanine at position 508 of NBD1, F508del, is found in ~90% of CF patients on at least one allele, defining the most common mutation causing CF. Impaired folding of F508del CFTR elicits its near-complete disposal by ER quality control mechanisms, and results in severe CF due to a marked reduction in apical membrane channel density. Significant amounts of WT CFTR are also degraded by most cells (Ward *et al.,* 1995), highlighting the complex folding landscape that even WT CFTR must traverse.

Approximately 2000 mutations of the CFTR gene, many quite rare, have been proposed as CF disease causing, while correction of the folding defect of F508del CFTR provides the greatest potential for enhancing the quality of life and life expectancy of CF patients. To date, the discovery of a small molecule, VX-809 (lumakaftor), which corrects 10-15% of F508del CFTR function *in vitro* (Van Goor *et al.,* 2011), has provided only limited improvement in clinical studies (Clancy *et al.,* 2012). The combination of lumakaftor with the gating potentiator, VX-770 (ivakaftor), has shown the potential to benefit a significant number of F508del patients, and their combination (Orkambi) has been approved for F508del homozygotes (Konstan *et al.,* 2017). Nevertheless, efforts to uncover the checkpoints in CFTR biogenesis where most F508del CFTR is lost to degradation pathways can be expected to identify targets whose modulation would further improve efficacy.

CFTR biogenesis is surveyed by molecular chaperones that monitor the protein’s conformational state. The core chaperone systems, Hsp70, Hsp90 and the Hsp40 co-chaperones, limit CFTR aggregation to facilitate its productive folding (Strickland *et al.,* 1997; Meacham *et al.,* 1999); however, these chaperones can also target CFTR mutants for degradation when the native fold is not achieved (Cyr *et al.,* 2002; Amaral, 2004). Unstable conformations of CFTR remain bound to chaperones, e.g. a prolonged association with Hsp70/Hsp90 recruits the ubiquitin ligase complex, Ubc5-CHiP, resulting in CFTR ubiquitylation and degradation by the 26S proteasome (Jensen *et al.,* 1995; Ward *et al.,* 1995; Meacham *et al.,* 2001; Sun *et al.,* 2006; Younger *et al.,* 2006). In contrast to the ATP-dependent core chaperone systems, small heat shock proteins (sHsps) represent a class of ATP-independent protein stabilizers. Generally, sHsps interact with intermediate, foldable protein conformations, e.g. during cell stress, and their clients are often refolded in association with ATP-dependent chaperones once stress conditions are relieved (Rajaraman *et al.,* 1996; Treweek *et al.,* 2000; Lindner *et al.,* 2001).

We recently identified a new sHsp-mediated pathway that promotes the selective degradation of F508del CFTR, while having little effect on the wild type (WT) protein (Ahner *et al.,* 2013). This pathway is initiated by binding of the small heat shock protein, Hsp27, to the F508del mutant, which recruits the SUMO E2 enzyme, Ubc9, to bring about the post-translational modification (PTM) of F508del CFTR by the <u>s</u>mall <u>u</u>biquitin-like <u>mo</u>difier, SUMO. Five SUMO paralogs are now considered to be functionally significant in mammalian systems, SUMO-1 through -5 (Owerbach *et al.,* 2005). This work will focus on SUMO-1 and SUMO-2/3. The latter two paralogs differ by only four amino acids and generally behave in a functionally similar manner, whereas SUMO-2/3 shares only 44% homology with SUMO-1. These SUMO paralogs are attached to their targets by isopeptide bonds, generally to lysine residues within the consensus motif ψKXD/E, where ψ is a hydrophobic residue, X is any amino acid and the modified lysine is followed by an acidic residue, a phosphorylated site, or a larger electronegative region. A strong physical interaction between Hsp27 and Ubc9, the single known E2 enzyme for SUMO conjugation, elicits primarily SUMO-2/3 modification of the CFTR mutant. The SUMO-2/3 paralogs contain a consensus sequence for SUMOylation, allowing them to form SUMO poly-chains. These chains are recognized by the SUMO-targeted ubiquitin ligase (STUbL), RNF4, which then ubiquitylates F508del CFTR via its RING domain, targeting it for degradation by the ubiquitin-proteasome system, UPS (Ahner *et al.,* 2013).

As a post-translational modification, SUMOylation impacts proteins that participate in diverse cellular processes, including nuclear transport, the maintenance of genome integrity, transcriptional control and the regulation of innate immune signaling (Adorisio *et al.,* 2017). SUMO modification is present in several neurodegenerative diseases that arise from the aggregation of misfolded proteins. Importantly, this PTM can influence the stability and solubility of misfolded and/or aggregated proteins (Liebelt and Vertegaal, 2016).

The SUMOylation cascade resembles that for ubiquitin modification, consisting of a series of thiol-mediated transfer reactions, but these pathways differ in that the single-known SUMO E2, Ubc9, can transfer SUMO directly to its client protein without intervention of an E3. The principal functions of E3 enzymes in the SUMO pathway appear to be assistance with target protein selection and catalytic transfer of the modifier (Gareau and Lima, 2010). The SUMO E3 ligases include members of the <u>p</u>rotein <u>i</u>nhibitor of <u>a</u>ctivated STAT (PAIS) proteins, an important family in signal transduction, which includes PIAS1, PIAS2 (also termed PIASx with isoforms PIASxα and PIASxβ), PIAS3 and PIAS4 (aka PIASy). Among the family members, PIAS4 has received attention due to its participation in signaling pathways that participate in oncogenesis and immune system regulation (Shuai and Liu, 2005; Rytinki *et al.*, 2009).

In the present work, we examined the influence of PIAS4 on WT and F508del CFTR, motivated by its identification when our protein array was probed with poly-SUMO-3, which had also attracted our attention to the STUbL, RNF4 (Ahner *et al.*, 2013). The identification of PIAS4 and the implication that it bound SUMO poly-chains suggested the hypothesis that it would support or enhance the Hsp27-Ubc9 mediated F508del CFTR degradation pathway described previously (Ahner *et al.*, 2013). In contrast, PIAS4 expression in CFBE airway cell lines, transduced to stably express either F508del or WT CFTR, promoted CFTR biogenesis rather than degradation, and therefore, we pursued its mechanism of action.

## RESULTS

### PIAS4 enhances CFTR biogenesis and F508del corrector efficacy

First, we examined the impact of over-expressing PIAS4 on CFTR protein levels. CFBE41o-airway epithelial cells stably expressing wild-type (WT) or F508del CFTR (hereafter, CFBE-WT or CFBE-F508del) were transfected with constructs to express PIAS4 or empty vector as control, followed by immunoblot (IB) for CFTR and PIAS4. As illustrated in Fig. 1A, PIAS4 over-expression augmented the steady-state expression level of immature (band B) and mature (band C) CFTR; the increases in WT and F508del levels were ~3-fold for band B and ~2-fold for band C on average, as shown in the accompanying quantitation of the aggregate data. The relative increases in band B exceeded those in band C, i.e. PIAS4 decreased the C/B band ratio. These findings suggest that the primary effect of PIAS4 is on the production and/or stability of the immature, ER-localized form of the WT and mutant proteins, a point to which we will return. Since the transfection efficiency of CFBE cells varies between 30-50%, the observed ~3-fold increase in band B is likely 2-3 times smaller than would result if the entire cell population were expressing PIAS4.

**Figure 1.**
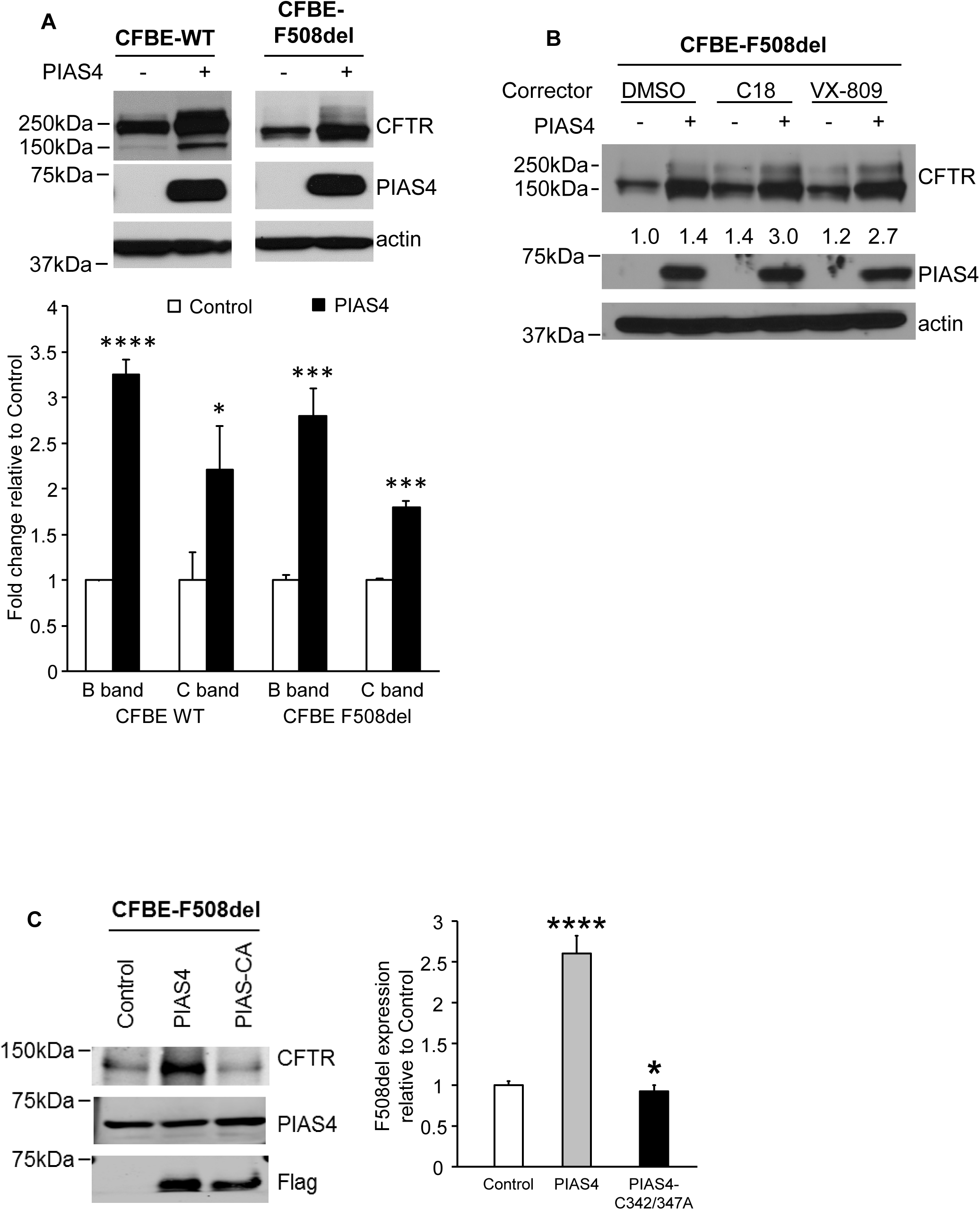
PIAS4 increases CFTR expression in the CFBE airway cells. (A) Over-expression of PIAS4 increases the steady-state expression of WT and F508del CFTR. Stable CFBE-WT and -F508del cells were transiently transfected with or without Flag-PIAS4 as described in Materials and Methods. Whole cell lysates were extracted 48 h after transfection and protein expression detected by immunoblot (IB) using the indicated antibodies. CFTR protein levels were quantified and normalized to control from three independent experiments. (*, p = 0.014; ***, p < 0.001; ****, p < 0.0001). (B) PIAS4 enhances the efficacy of CFTR correctors in CFBE cells stably expressing F508del CFTR. Flag-PIAS4 was expressed in CFBE-F508del cells as described in A. After 24 h, the transfected cells were treated with DMSO, 5 μM C18 or 5 μM VX-809 for 24 h, then cells were lysed and analyzed by IB. The numbers below the CFTR blots give the band C densities relative to control (DMSO) (C) The impact of PIAS4 on CFTR expression depends on its SUMO E3 ligase activity. Flag-PIAS4 and its catalytic mutant, Flag-PIAS4-CA, were transiently transfected into CFBE-F508del cells. Whole cell lysates were examined by IB using the indicated antibodies. CFTR signals were normalized to control values in three independent experiments (*, p < 0.05; ****, p < 0.0001).

Second, we determined whether the augmented level of immature F508del CFTR would enhance the ability of correctors to generate the mature form of the mutant protein.

Experiments similar to those of Fig. 1A were performed, in which the CFBE-F508del cells were treated for 24 hrs with either vehicle, 5 μM VX-809 (Van Goor *et al.,* 2011) or one of its developmental precursors from the same chemical series, 5 μM C18 (www.cftrfoldinq.orq). As shown in Fig. 1B, the expression of PIAS4 alone elicited a marked increase in immature F508del CFTR, together with a small increase in mature protein, noted also in Fig. 1A. However, a more substantial level of F508del band C was found when PIAS4 over-expression was combined with either of the correctors. The band C densities relative to control, shown below the F508del bands, indicate that PIAS4 was as effective as the correctors at increasing the mature mutant protein level, and when they were combined, PIAS4 increased corrector efficacy ~2-fold.

Third, we performed a negative control experiment to determine whether PIAS4 was acting as a SUMO E3; that is, to assess the requirement for the catalytic cysteines of the SP-RING domain of PIAS4 in its action on F508del expression. The experiments were performed as in Fig. 1A and the results shown in Fig. 1C: mutation of its two catalytic cysteines (C342 and C347) to alanine eliminated the PIAS4-induced increase in F508del CFTR expression. In fact, although the difference is small, expression of the PIAS4-CA mutant significantly reduced F508del expression level below that of the control. This finding suggests that the mutant interfered with the action of endogenous PIAS4, consistent with the possibility that this SUMO E3 makes a physiological contribution to CFTR biogenesis.

The latter concept was further evaluated in PIAS4 knockdown experiments. Stable CFBEWT or -F508del cell lines were transduced with shRNA targeting PIAS4 or with a scrambled shRNA control; these constructs were expressed using an adenovirus vector, described in Materials and Methods. After 72 hrs, cell lysates were prepared for CFTR IB, and the results and mean data are illustrated in Fig. 2A&B. Knockdown of the SUMO E3 was very effective, and produced reductions in the expression level of WT and F508del CFTR that averaged ~40%. These findings, together with those of Fig. 1C, further support the conclusion that endogenous PIAS4 plays a role in promoting the biogenesis of WT and F508del CFTR.

**Figure 2.**
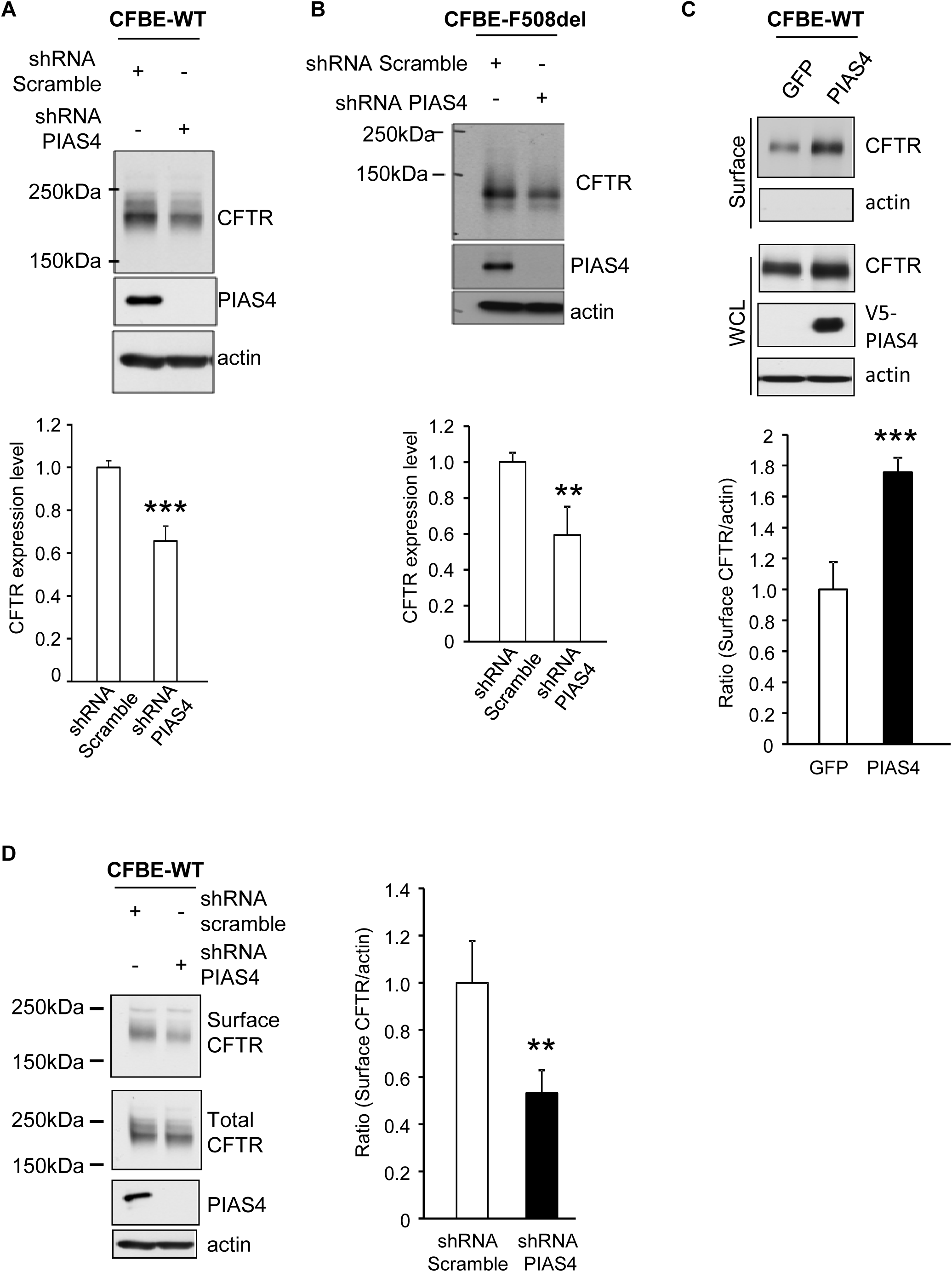
PIAS4 knockdown decreases CFTR expression in CFBE airway cells. (A&B) Knockdown of PIAS4 reduces CFTR expression in stable CFBE-WT and -F508del cells. shRNA targeting PIAS4 was transduced using a recombinant adenovirus, and compared with a scrambled shRNA. CFTR expression was evaluated by IB with the indicated antibodies after 48 h exposure to virus. For all experiments in Fig. 2, bar graphs provide composite means ± SEM from at least four independent experiments (**, p < 0.01; ***, p < 0.001). (C) Over-expression of PIAS4 promotes CFTR cell surface expression. CFBE-WT cells were transduced with PIAS4 or GFP for 48 h and biotinylation assays performed. Streptavidin elution was followed by IB with the indicated antibodies, as described in Materials and Methods. Surface CFTR signals were normalized to the actin loading control. (D) PIAS4 knockdown reduces CFTR surface expression. CFBE-WT cells were transduced with adenovirus vector shRNA targeting PIAS4 or scrambled shRNA for 48 h and then subjected to surface biotinylation and analysis as described in part C and Materials and Methods.

Next, we used cell surface biotinylation to evaluate the impact of PIAS4 over-expression or knockdown on CFTR expression in the airway cell plasma membrane (PM). For these studies, CFBE cells stably expressing WT CFTR were transduced to express either PIAS4 or GFP, then the cell surface was biotinylated as described in Materials and Methods. Cell lysates were prepared and biotinylated PM proteins were pulled down using streptavidin agarose, IBs were performed for CFTR and actin, and the cell lysates were blotted for the indicated proteins. As shown in Fig. 2C, the over-expression of PIAS4 increased cell surface CFTR ~ 75% on average. Actin was not detected in the pulldown, a negative control indicating that labeling was restricted to proteins resident at the cell surface. Thus, the increase in WT CFTR levels in cells over-expressing PIAS4 leads also to increased CFTR density in the plasma membrane, which should lead to an increase in CFTR-mediated anion transport; that data will be shown below.

Finally, the impact of reduced PIAS4 levels on the surface expression of WT CFTR was examined by performing PIAS4 knockdown followed by cell surface biotinylation. CFBE-WT cells were transduced to express PIAS4 or control shRNA and then subjected to surface biotinylation and streptavidin pulldown after 72 hrs; the results are illustrated in Fig. 2D. Reduced PIAS4 resulted in a ~ 50% reduction in cell surface CFTR protein, which paralleled the effects of reduced PIAS4 expression on CFTR expression levels (Fig 2A&B). Together, the data of Fig. 2, indicate that PIAS4 plays a significant role in determining the steady-state expression level of immature CFTR, as well as that of the mature WT protein that resides in the plasma membrane.

### PIAS4 slows the degradation of F508del CFTR in the ER

The marked increase in immature CFTR expression, observed for both WT and F508del, suggested that PIAS4 augments the production or stability of CFTR in the endoplasmic reticulum, where the immature forms of these proteins are localized (Gregory *et al.*, 1991; Ward and Kopito, 1994). To examine this hypothesis, we performed cycloheximide (CHX) chase experiments using CFBE-WT or CFBE-F508del stable cell lines that were transfected with constructs for PIAS4 or the vector control. Fig. 3A shows the time course of immature F508del CFTR expression once protein synthesis had been interrupted in PIAS4 or control cells. At the indicated times, cells were lysed and the lysates prepared for IB with the indicated antibodies. The data demonstrate that band B F508del CFTR disappeared with roughly exponential kinetics under control conditions with a t_1/2_ of ~1hr, as observed in previous studies (Liang *et al.*, 2012), whereas the t_1/2_ for disposal of the mutant in the PIAS4 over-expressing cells was ~3 hrs. The degradation of F508del under control conditions was mediated primarily by the proteasome, as reflected by its inhibition by MG132 (data not shown), findings that are consistent with prior studies (Ward and Kopito, 1994; Sun *et al.*, 2008; Liang *et al.*, 2012).

Next, we used the CHX chase protocol to evaluate the influence of PIAS4 over-expression on the kinetics of WT CFTR degradation, as shown in Fig. 3B. The pattern of immature CFTR disappearance was different from that observed for F508del in that PIAS4 elicited a somewhat more rapid decrease in immature WT CFTR, but did not stabilize the protein, as was observed for F508del band B. Rather, these data suggest that the stabilization of WT CFTR occurred primarily at the PM. Mature WT CFTR decreased to ~50% of its initial value at 4 hrs in control cells, but was more stable with PIAS4 over-expression, decreasing by only ~10% during the same period. A likely explanation for the small reduction in WT band B with increased PIAS4 relates to the impact of the SUMO E3 on band C, since the mature protein draws on the pool of immature CFTR in the ER. Thus, band B starts at a higher level with PIAS4 expression, but once protein synthesis is blocked by CHX, the high level of a more stable band C provides a ‘sink’ for immature WT CFTR remaining in the ER. Nevertheless, after several hours of CHX inhibition, PM CFTR begins to decline. Whether and how PIAS4 acts to increase CFTR stability following its exit from the ER will require additional studies, but increased peripheral stability would be expected to contribute to enhanced corrector action as shown in Fig. 1B.

**Figure 3.**
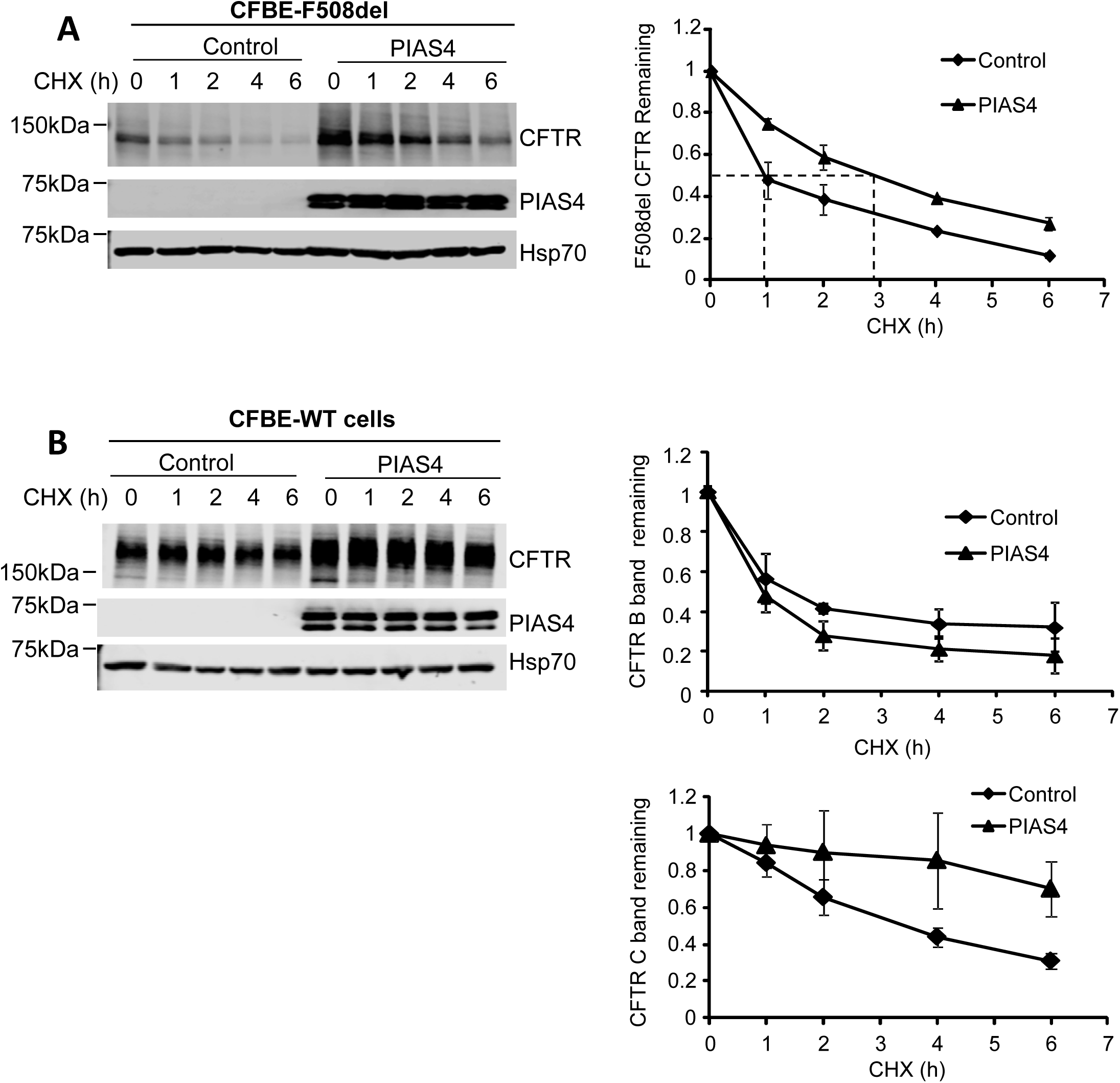
PIAS4 stabilizes CFTR, reducing its proteasomal degradation. (A) PIAS4 slows F508del CFTR degradation. CFBE-F508del cells were transfected with or without Flag-PIAS4 as described in Materials and Methods. After 48 h, the transfected cells were treated with 100 µg/ml CHX and lysed with RIPA buffer at the indicated CHX treatment times (0, 1, 2, 4, 6 h) to follow F508del (band B) expression during inhibition of new protein synthesis. Following IB, blot densities from four independent experiments were averaged to construct the relations for F508del CFTR (band B) remaining as a function of time, with values from each experiment normalized to the mean densities at the beginning of the CHX chase (t = 0). (B) PIAS4 stabilizes mature WT CFTR. Experiments performed as described in A, but with CFBE-WT cells. Time courses for expression of CFTR bands B and C relative to control are provided. See text for discussion.

### PIAS4 increases functional F508del CFTR at the cell surface

To determine whether the increase in immature F508del expression elicited by PIAS4 translates into increased trafficking to the plasma membrane during corrector treatment, we examined the cell surface expression of fluorogen activating protein (FAP)-tagged F508del CFTR in CFBE cells treated with VX-809. The FAP labeling approach has been used previously to compare different correctors for their ability to mobilize FAP-tagged F508del CFTR to the PM (Holleran *et al.*, 2012) or to monitor its intracellular trafficking itinerary following its endocytic retrieval from the PM (Holleran *et al.*, 2013). CFBE cells stably expressing FAP-F508del CFTR were transferred to a 96-well plate with or without recombinant adenovirus encoding PIAS4. After 24 hrs, the cells were incubated with 2 μM VX-809 and read for FAP activation the following day. The panels of Fig. 4A show cell surface labeling of FAP-F508del in the absence and presence of PIAS4 co-expression, respectively. The accompanying bar graph, Fig. 4B, provides the mean surface intensity, which was increased ~ 2.5-fold by PIAS4, in agreement with the biochemical data that reported a 2.7-fold increase in band C of F508del (Fig. 1B).

**Figure 4.**
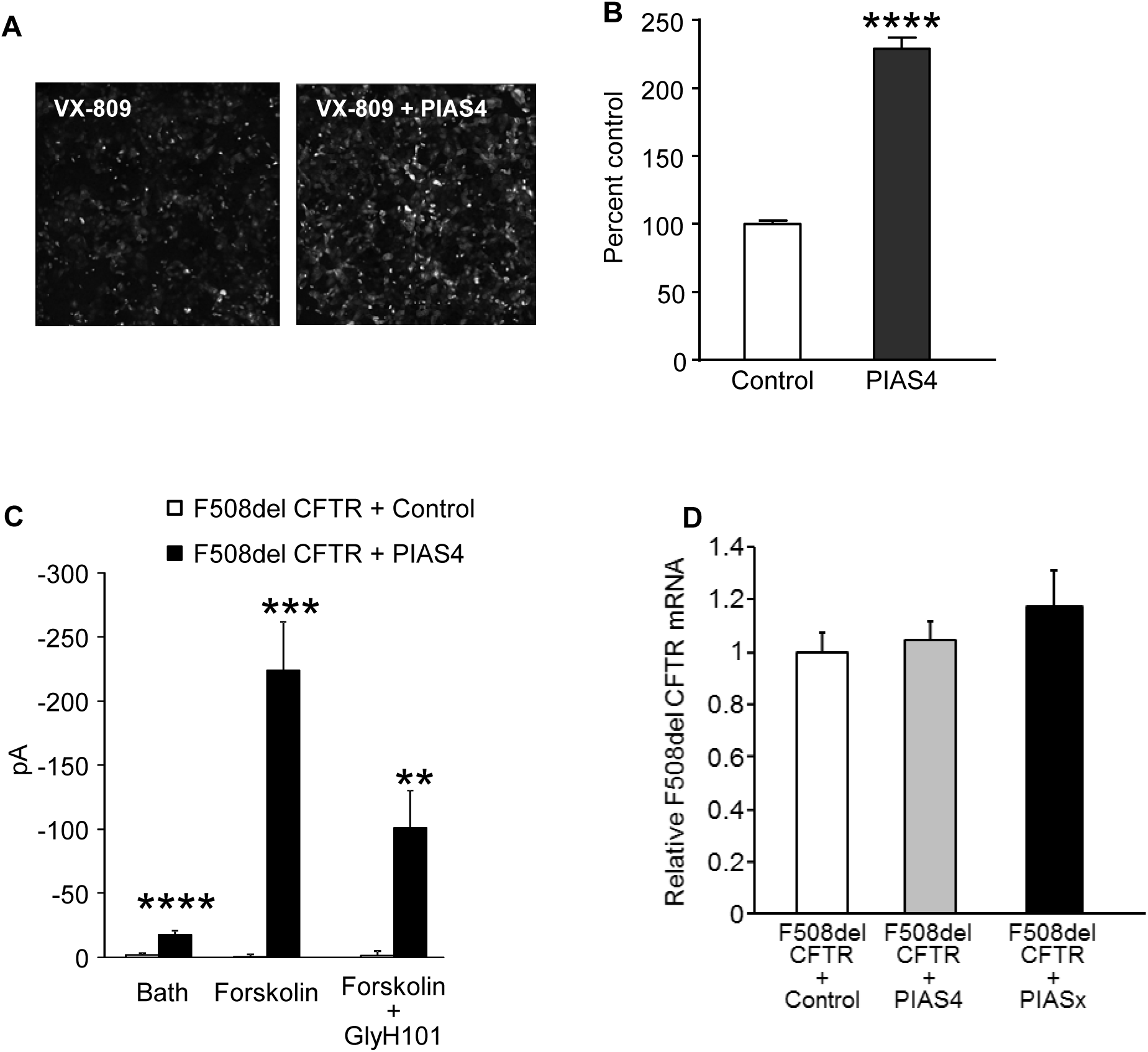
PIAS4 increases functional F508del CFTR at the surface of airway cells. (A) The membrane-impermeant fluorescent dye, MG-B-Tau, was used to detect FAP-F508del CFTR in CFBE-F508del cells stably expressing the mutant CFTR and treated with 2 µM VX-809 overnight. Fluorescence imaging reveals an increase in FAP-F508del CFTR surface expression when the VX-809 treated CFBE cells were transduced with recombinant adenovirus expressing PIAS4. Representative images from four independent experiments are shown. (B) The bar graph represents the quantified relative surface expression of FAP-F508del CFTR in CFBE cells in the presence of 2 µM VX-809 under control conditions or in CFBE cells transduced with recombinant adenovirus expressing PIAS4. The experiment was repeated four times (****, p< 0.0001). (C) Chloride currents in PIAS4 expressing CFBE41o- parental airway cells. Whole-cell patch clamp was used to monitor baseline Cl currents and the effects of forskolin and the inhibitor, GlyH-101. Control cells were transfected with F508del CFTR and EGFP; experimental cells transfected with F508del CFTR plus PIAS4 and EGFP. The fluorophore was used to identify expressing cells. The pipette and bath solutions contained equal concentrations of Cl and currents were measured at a holding potential of -40 mV (**, p = 0.006; ***, p = 0.0003; ****, p < 0.0001). (D) The impact of PIAS4 on F508del CFTR expression is not detected at the mRNA level. Three constructs: empty vector (control), Flag-PIAS4 or Flag-PIAS2 were transfected into CFBE-F508del stable cells. After 48 h, total RNA was extracted and subjected to qPCR as described in Materials and Methods. β-actin was used as an internal control.

We also examined the functionality of cell surface F508del CFTR in parental CFBE41o- cells 2 days following transfection with F508del CFTR using patch clamp methods to record their whole-cell currents, as indicted in Materials and Methods. The pipette and bath solutions contained equal Cl concentrations and the forskolin-stimulated currents, recorded at a holding potential of -40 mV, were inhibited by GlyH-101 (Muanprasat *et al.*, 2004). These findings indicate that mutant CFTR that traffics to the PM exhibits agonist stimulated, CFTR-mediated, Cl conductance properties.

In principle, PIAS4 might augment CFTR biogenesis by increasing CFTR transcription, translation or by its post-translational modification. To rule out a change in F508del CFTR message levels, we evaluated the influence of PIAS4 expression on CFTR mRNA levels by qPCR; the results are shown in Fig. 4D. The levels of mRNA from CFBE-F508del cells did not differ between cells transfected with empty vector, PIAS4 or the PIAS2 isoform (aka PIASx); the latter also had no effect the expression level of CFTR (data not shown).

### PIAS4 drives SUMO-1 conjugation of CFTR and reduces its SUMO-2 modification

Our prior work demonstrated that the small heat shock protein, Hsp27, primarily interacted with F508del CFTR and associated with the SUMO E2 enzyme, Ubc9, to modify F508del CFTR with SUMO-2/3 (Ahner *et al.*, 2013; Gong *et al.*, 2015). This Hsp27/Ubc9 pathway led to the formation of SUMO poly-chains and their recognition by the SUMO-targeted ubiquitin ligase, RNF4, promoting mutant CFTR degradation by the proteasome (Ahner *et al.*, 2013). However, the stabilizing effect of PIAS4 on WT and F508del CFTR suggested that an alternative pathway leads to the opposing outcome in response to intervention of the SUMO E3, which supports stabilization and biogenesis of the WT and mutant proteins, rather than degradation.

The hypothesis that a different SUMO paralog modifies CFTR consequent to its interaction with PIAS4 was evaluated initially using *in vitro* SUMOylation experiments performed with purified SUMO pathway components and human F508del NBD1. In a previous study, we found that purified F508del NBD1 was preferentially conjugated to SUMO-2/3 when Hsp27 was introduced into the reaction mixture *in vitro* (Gong *et al.*, 2015). This outcome was similar to the SUMO-2 modification of full-length F508del CFTR observed during Hsp27 over-expression *in vivo* (Ahner *et al.*, 2013). As shown in Fig. 5A, F508del NBD1 could be modified by SUMO-1 or -2 *in vitro;* its conjugation with SUMO-2 was somewhat greater than that by SUMO-1, as found previously (Gong *et al.*, 2015). However, the addition of purified PIAS4 to the reaction mix led to increased F508del NBD1 conjugation with SUMO-1, plus a smaller but significant decrease in its modification by SUMO-2. Quantitation of the changes from all experiments showed that PIAS4 induced a ~80% increase in SUMO-1 modification of the mutant NBD1 and a ~30% decrease in its conjugation with SUMO-2.

**Figure 5.**
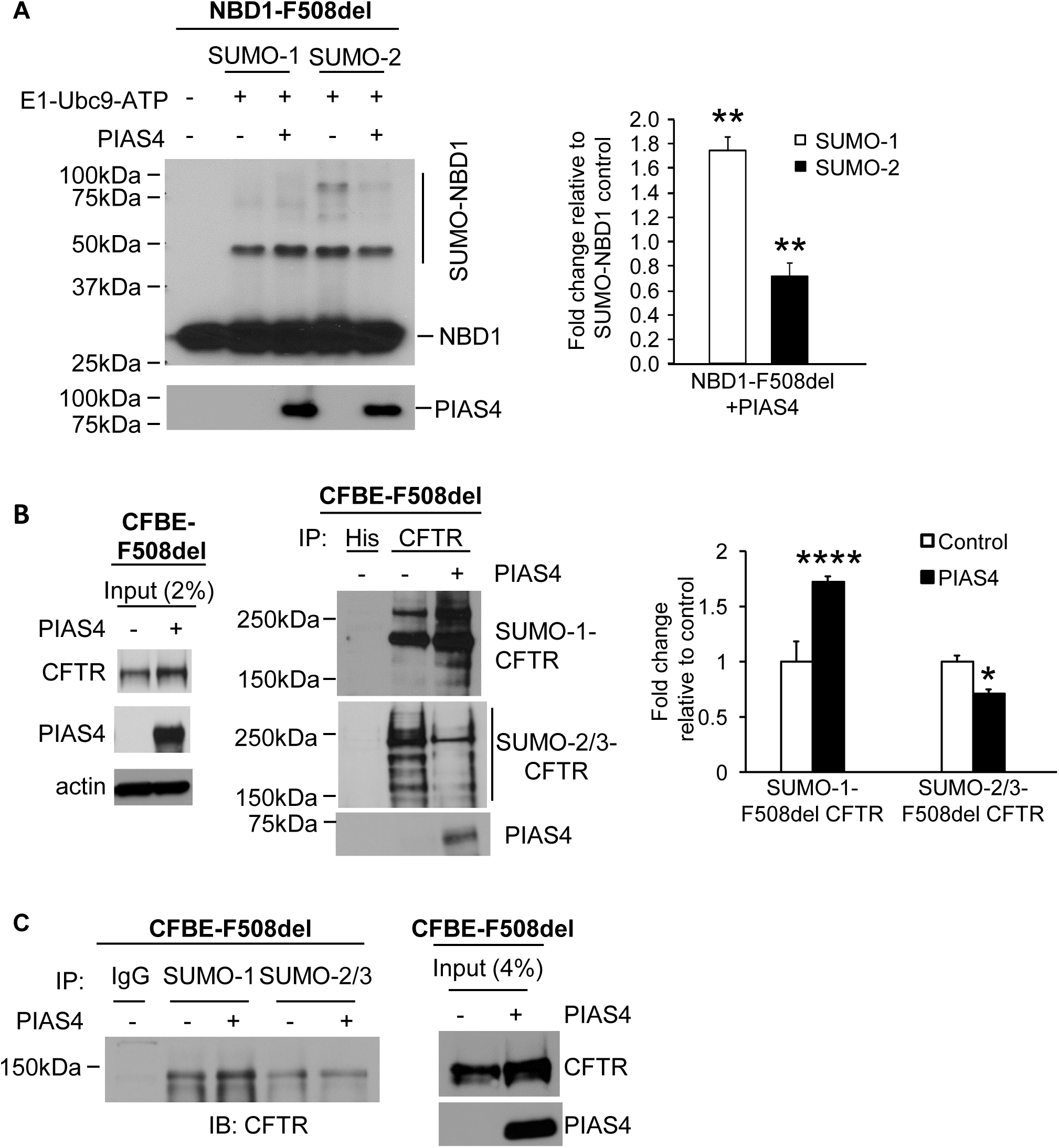
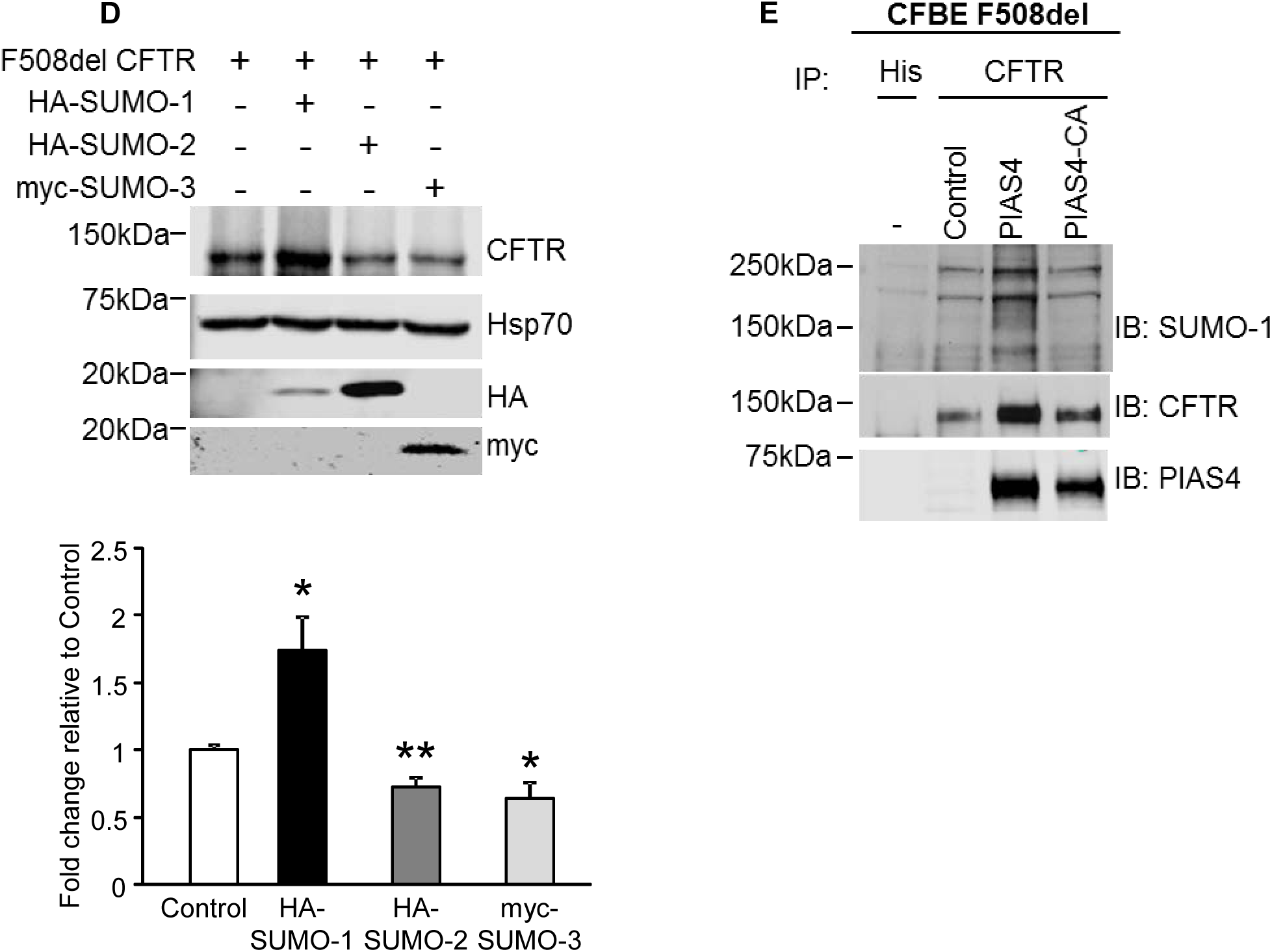
PIAS4 facilitates SUMO-1 conjugation to F508del CFTR and reduces its modification by SUMO-2/3. (A) PIAS4 promotes the conjugation of NBD1-F508del to SUMO-1 *in vitro* and reduces that by SUMO-2/3. Purified F508del-NBD1-1S (Rabeh *et al.,* 2012) was incubated with purified SUMOylation components for and shown previously (Gong *et al.,* 2016). The reaction mixture was resolved on 12% SDS-PAGE and NBD1 was detected by anti-CFTR-NBD1 (#660). The modified NBD1-F508del-1S protein signals were quantified and normalized to control values from three independent experiments. (**, p < 0.01). (B) *In vivo,* PIAS4 induces F508del CFTR conjugation to SUMO-1, while reducing its modification by SUMO-2/3. Flag-PIAS4 was over-expressed in CFBE-F508del cells; empty vector as control. CFTR was immunoprecipitated (IP’d) from cell lysates and immunoblotted (IB’d) with antibodies against endogenous SUMO paralogs. The SUMO signals were normalized to control values in three independent experiments (*, p = 0.046; ****, p = 0.00006). (C) PIAS4 impacts the interactions of F508del CFTR with endogenous SUMO isoforms in CFBE-F508del cells. Co-IP experiments performed as in (B) using lysates from cells with and without PIAS4 over-expression. SUMO paralogs interacting with F508del CFTR were detected in the IP using anti-SUMO-1 or -2/3 with rabbit IgG as control. The experiment was repeated three times with equivalent results. (D) PIAS4 actions on F508del CFTR biogenesis or degradation were reproduced by expression of active SUMO paralogs that express the C-terminal di-glycine (G-G) motif, whose exposure by SUMO peptidase is required prior to target conjugation (Pichler *et al.,* 2017). Parental CFBE41o- cells were transfected with F508del CFTR plus HA-SUMO-1 or -2 or myc-tagged SUMO-3, and after 48 h, the expression levels of F508del CFTR were determined by IB. The bar graph illustrates the composite data, which shows that activated SUMO-1 expression reproduces the actions of PIAS4 on F508del biogenesis and that of SUMO-2/3 mimics RNF4 to promote F508del degradation, data from four experiments relative to control (*, p < 0.05; **, p <0.01). (E) The SUMO E3 ligase activity of PIAS4 is required for its facilitation of F508del CFTR modification by SUMO-1. CFBE-F508del cells were transiently transfected with vector, Flag-PIAS4 or the RING domain catalytic mutant, Flag-PIAS4-CA. Cell lysates were (IP’d) with anti-CFTR followed by (IB) with the indicated antibodies for SUMO-1, CFTR or PIAS4.

Next, we asked whether differential paralog modifications of full-length F508del CFTR *in vivo* are also influenced by over-expression of PIAS4; the results are shown in Fig. 5B. CFBE-F508del cells were transfected with PIAS4 or the corresponding empty vector. After immunoprecipitation (IP) of CFTR, the pulldowns were blotted for either SUMO-1 or SUMO-2/3. The input blots in the left panel show the increase in F508del expression observed with co-expression of PIAS4, as in prior figures. Following IP of mutant full-length CFTR, the SUMO paralog IBs indicate that F508del CFTR was primarily modified by SUMO-1 during PIAS4 co-expression, and this was accompanied by a reduction in modification of the mutant by SUMO-2/3.

Qualitatively similar results to those shown in Fig. 5B were obtained when the experiment was performed using the same conditions but in the opposite manner, i.e. SUMO IP followed by F508del CFTR IB (Fig. 5C). The IP of SUMO-1 pulled down more F508del CFTR from cells co-expressing PIAS4 than observed for the vector control, whereas the IP of SUMO-2/3 did not isolate more F508del CFTR from PIAS4 expressing cells, despite the increased levels of the CFTR mutant (see input).

While the experiments in Figs. 5B&C relied on endogenous SUMO paralogs to examine their differential effects on F508del CFTR fate, similar patterns could also be produced by expressing pre-activated paralogs that expose the C-terminal G-G motif that is required prior to conjugation. The parental CFBE41o- cell line was transfected with F508del CFTR and epitope-tagged active SUMO paralogs, and their impact on the expression levels of the mutant protein determined after one day by IB. As illustrated in Fig. 5D, the expression of active SUMO-1 elicited a ~75% increase in F508del band B relative to control, thus mimicking the biogenic effect seen with PIAS4 over-expression. Contrasting with this were the effects of active SUMO-2 and -3, which produced decreases of ~30-40% in expression of immature F508del vs. control, paralleling the actions of Hsp27/RNF4 in promoting mutant protein degradation.

Finally, to verify that SUMOylation of F508del CFTR could be attributed to the E3 catalytic activity of PIAS4, its ability to SUMO-1 modify F508del CFTR was assessed using mutant PIAS4 in which the catalytic cysteines of the SP-RING domain were mutated to alanine. As seen in Fig. 5E, and previously in Fig. 1C, these C/A mutations eliminated the ability of PIAS4 to increase F508del expression level. This finding is consistent with reduction in mutant CFTR pulled down in the IP relative to that with WT PIAS4; in addition, this protein was much less modified by SUMO-1 than that for the catalytically active SUMO E3. Together, these findings indicate that PIAS4 can alter the balance between post-translational SUMO paralog modifications of F508del NBD1.

### PIAS4 reduces F508del ubiquitylation and the impact of RNF4

The above findings indicate that PIAS4 modifies FL F508del CFTR *in vivo* similarly to that of F508del NBD1 *in vitro*, increasing its conjugation with SUMO-1. The modification of FL F508del or NBD1 by SUMO-1 was associated also with a reduction in the conjugation of mutant CFTR with SUMO-2/3 (Fig. 5 A&B). Since this PTM can lead to formation of SUMO poly-chains and interaction with the SUMO-targeted ubiquitin ligase (STUbL), RNF4, a reduction in ubiquitin modification of the mutant would be anticipated.

The results of this experiment are shown in Fig. 6A. CFBE-F508del stable airway cells were transfected with either PIAS4 or empty vector as control. Both PIAS4 and control cells were incubated with proteasome inhibitor MG132 for 6 hrs to block rapid degradation of the ubiquitylated F508del CFTR which would otherwise occur. Following F508del CFTR IP, the pull downs were blotted for CFTR and ubiquitin, revealing an increased CFTR level in the IP as well as a ~50% reduction in the level of F508del ubiquitylation in response to PIAS4 over-expression; this effect was variable, as reflected by the SEM that characterizes the aggregate data.

**Figure 6.**
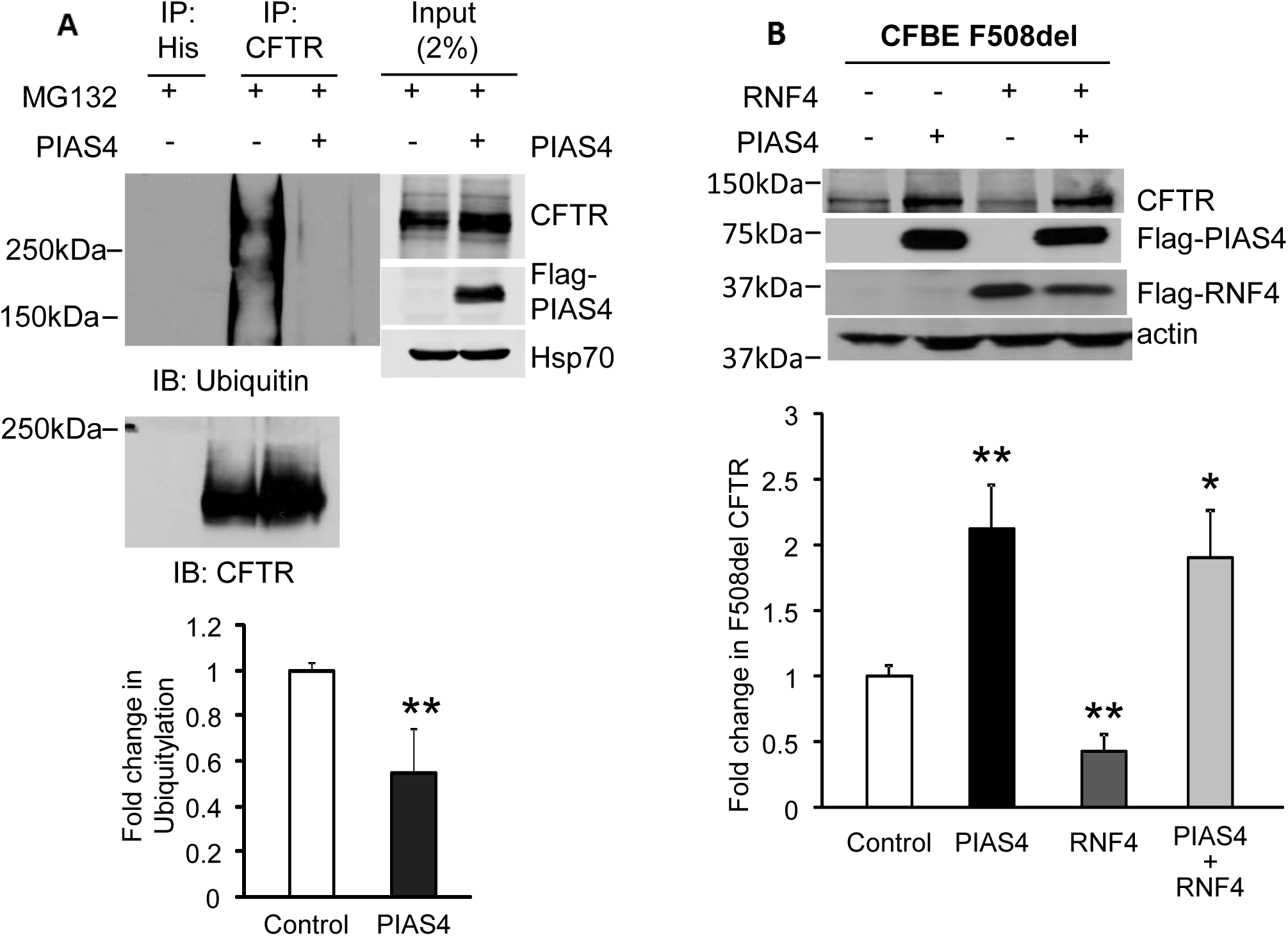
PIAS4 protects F508del CFTR from proteasomal degradation. (A) PIAS4 expression decreased the ubiquitylation of F508del CFTR. CFBE-F508del stable cells were transfected with Flag-PIAS4 or empty vector. After 48 h, cells were treated with 50 µM MG132 for 5 h. Cell lysates were immunoprecipitated with CFTR antibody and the IPs probed for ubiquitin by IB. The ubiquitin signals were normalized to control values from three independent experiments (**, p = 0.003). (B) PIAS4 obviates RNF4-mediated F508del CFTR degradation. Flag-PIAS4 and empty vector or Flag-RNF4, were over-expressed in CFBE-F508del cells. After 48 h, cells were lysed and subjected to IB with the indicated antibodies. The CFTR signals were quantified and normalized to control from five independent experiments. (*, p = 0.01; **, p = 0.005 or 0.002).

In principle, the effects of PIAS4 and RNF4 would antagonize one another since RNF4 increases F508del ubiquitylation (Ahner *et al.*, 2013), whereas PIAS4 reduced F508del ubiquitin modification (Fig. 5A). Therefore, we asked whether the decrease in F508del CFTR expression induced by RNF4 co-expression could be offset by over-expression of PIAS4. CFBE-F508del stable cells were transfected with either PIAS4, RNF4, or their combination, and the steady-state levels of F508del CFTR were determined after 48 h. The composite results are provided in Fig. 6B. PIAS4 over-expression increased immature F508del CFTR levels 2.1-fold, while expression of RNF4 reduced F508del band B levels ~60%. When these post-translational modifiers were combined, PIAS4 reversed the pro-degradation impact of RNF4. This outcome likely results from competition between the two SUMO paralogs that mediate the actions of PIAS4 and RNF4, with the impact of PIAS4 on mutant CFTR fate being more significant.

Finally, we examined the influence of four predicted SUMOylation sites (K377, K447, K1199 and K1468) on F508del CFTR expression levels following their mutation to arginine. These sites were predicted by the software, GPS-SUMO, found at: http://sumosp.biocuckoo.org. As shown in Fig. 7, PIAS4 increased the expression of F508del by nearly 4-fold in the positive control, but mutation of four predicted SUMOylation sites virtually eliminated the ability of PIAS4 to augment CFTR biogenesis. The rise in expression of the F508del CFTR-4KR mutant relative to the CFTR mutant lacking these site mutations most likely reflects suppression of the degradation pathway involving SUMO-2/3 and RNF4. These data are consistent with the hypothesis that the pathways directing biogenesis vs. degradation of CFTR by different SUMO paralogs are competitive, and that they likely target the same sites for SUMO modification.

**Figure 7.**
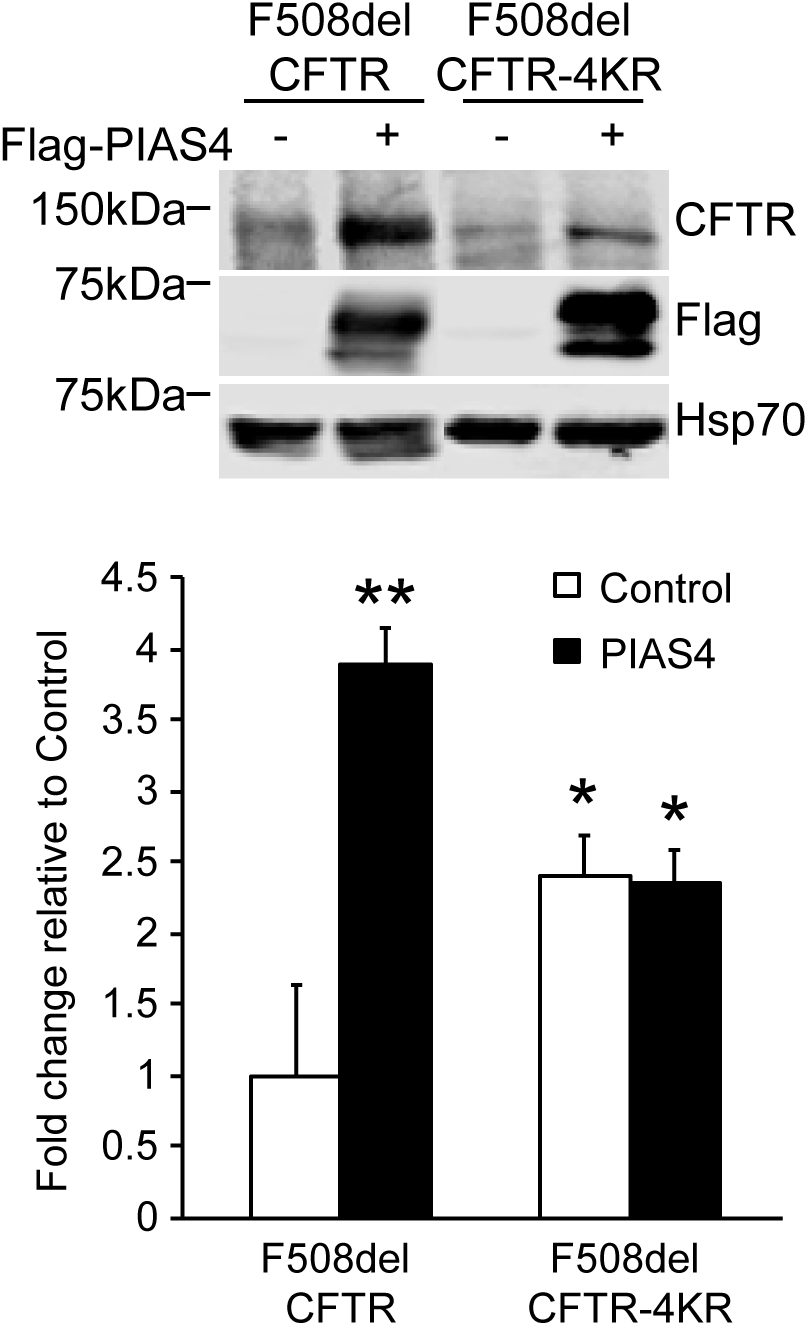
PIAS4 can stabilize F508del CFTR by modifying four consensus SUMOylation sites. F508del CFTR with K/R mutations at predicted SUMOylation sites (K377, 447, 1199, and 1468R) is refractory to the impact of PIAS4 on mutant CFTR biogenesis. F508del CFTR or F508del CFTR-4KR were transfected with Flag-PIAS4 or vector control into CFBE41o- parental airway cells. Whole cell lysates were extracted and protein expression levels detected by IB with the indicated antibodies. Data from four independent experiments (* p< 0.05; **, p = 0.01).

## DISCUSSION

SUMOylation targets a variety of cellular processes and has been implicated in numerous diseases, including diseases of protein misfolding (Gareau and Lima, 2010; Srikanth and Verma, 2017). SUMO modification occurs on transcription factors, membrane receptors, ion channels, etc. (Benson *et al.*, 2017). Tandem mass spectrometry has identified SUMOylation sites overlapping with other post-translational modifications such as ubiquitylation, acetylation and methylation. SUMO knockout is embryonic lethal, and it has been estimated that a significant fraction of the mammalian proteome is subject to SUMO modification (Hendriks and Vertegaal, 2016). Five SUMO paralogs have been described: SUMO-1 through SUMO-5. SUMO-4 can be difficult to conjugate to substrates due to a proline residue that restricts its isopeptidase activity (Owerbach *et al.*, 2005). It, and SUMO-5, have limited tissue distributions and have been studied less extensively that the other paralogs (Liang *et al.*, 2016). Sometimes referred to as the twins, SUMO-2 and -3 are 97% identical; and as no antibody can distinguish these paralogs, they are usually identified as SUMO-2/3. SUMO-1 has 46% identity with SUMO-2/3, and these three paralogs are widely expressed. SUMO modification is covalent and can be reversed by SUMO-specific peptidases.

### Hsp27/SUMO-2/3/RNF4 and F508del CFTR degradation

Numerous pathways are available for disposing of misfolded proteins. Core-glycosylated substrates that fail to fold properly such as the common CFTR mutant, F508del, undergo glucose- and mannose-trimming as a critical quality control mechanism. The lectin EDEM seems to be essential for the recognition and targeting of misfolded CFTR channels to ubiquitin mediated degradation (Gnann *et al.*, 2004). Subsequently, the ubiquitin E3, RMA1 (RNF5) mono-ubiquitylates F508del CFTR and gp78 mediated poly-ubiquitylation follows (Younger *et al.*, 2006; Morito *et al.*, 2008). SYVN1 (HRD1) has been shown to support RMA1/gp78 facilitated modification dependent on its catalytic activity (Ramachandran *et al.*, 2016). While RMA1 responds to early folding defects, late in the maturation process, after translation has concluded, Csp recruits another E3 ligase, CHiP, to promote CFTR ubiquitylation (Meacham *et al.*, 2001; Schmidt *et al.*, 2009). As discussed above, another PTM, Hsp27 mediated SUMO-2/3 modification targets F508del CFTR to RNF4 catalyzed ubiquitylation and proteasomal degradation (Ahner *et al.*, 2013). If these two pathways are completely independent and if they target different pools of folding mutants remains to be determined. Other ubiquitin E3 ligases implicated as having the capacity to catalyze F508del degradation include: SCF, NEDD4L, and RNF185 (Yoshida *et al.*, 2002; Caohuy *et al.*, 2009; El Khouri *et al.*, 2013). In some cases, the evidence for their involvement in misfolded CFTR disposal derives from their exogenous expression in model systems, and may not reflect their physiological significance in the cell types affected by CF.

Previously, we identified an interaction between F508del CFTR and the small heat shock protein, Hsp27, which cooperated with the SUMO E2 enzyme, Ubc9, to modify the CFTR mutant with SUMO-2/3 (15). The modification of F508del CFTR with SUMO-2/3 did not require the intervention of a SUMO E3 as the SUMO E2, Ubc9, can perform enzymatic conjugation of SUMO to target proteins. The poly-chains formed by SUMO-2/3 are recognized by RNF4, a SUMO-targeted ubiquitin ligase having four SUMO interacting motifs (SIMs) at its N-terminus (Tatham *et al.*, 2008), leading F508del CFTR to ubiquitylation and degradation by the UPS. Then we can ask: why would another ubiquitin E3 ligase be needed to degrade F508del CFTR in addition to the eight ubiquitin E3 pathways listed above? This question prompted us to explore identification of other SUMO binding proteins that might interact with WT or F508del CFTR using a protein array probed for binding of either mono-SUMO-1 or SUMO-3 poly-chains; this array had previously identified RNF4 and several SUMO pathway components. The SUMO E3, PIAS4 was a prominent candidate on that list, as it is known to have two SUMOylation sites and eight SUMO interacting motifs (SIMs).

### PIAS4/SUMO-1 and CFTR Biogenesis

The results of this study showed that PIAS4 stimulated the biogenesis of both WT and F508del CFTR, thus opposing the pro-degradative pathway comprised of Hsp27-Ubc9-SUMO-2/3-RNF4 (Ahner *et al.*, 2013). All experiments here were performed in CFBE airway cells, either the parental line, CFBE41o- (Bruscia *et al.*, 2002), which expresses little or no endogenous F508del CFTR, or using cells derived from this line and transduced to stably express WT or F508del CFTR (Bebok *et al.*, 2005). The action of PIAS4 on WT CFTR was somewhat unexpected since the Hsp27 pathway selectively targeted F508del degradation, with no significant activity against the wild-type protein. This behavior results from detection of misfolded F508del by the sHsp, which targets a transitional (partially denatured) conformation of the mutant (Gong *et al.*, 2016). For WT CFTR, PIAS4 increased most strongly the steady-state expression of immature CFTR (3.2-fold) while increasing the mature protein by 2.2-fold. Somewhat smaller but proportionately similar increases in F508del expression (2.8-fold for band B vs. 1.8-fold for band C, Fig. 1A) were produced by PIAS4. Moreover, PIAS4 augmented the response to the class I correctors C18 and VX-809 (Okiyoneda *et al.*, 2013) up to three-fold over control (Fig. 1B), and this was confirmed by cell surface biotinylation (Fig. 2C&D) and by surface labeling of FAP-tagged F508del CFTR (Fig. 4A). Currents generated by F508del CFTR that was rescued to the plasma membrane by PIAS4 were stimulated by forskolin and inhibited by GlyH-101, as detected using whole-cell patch clamp.

The physiological significance of PIAS4’s impact was reflected by its knockdown, which reduced the steady-state CFTR expression levels in CFBE-WT or -F508del cells (Fig. 2A&B), as well as decreasing the cell surface expression of WT CFTR (Fig. 2D). Conversely, over-expression of PIAS4 augmented the cell surface expression of WT CFTR nearly 2-fold (Fig. 2C). In cycloheximide chase experiments, the increase in immature (band B) F508del was traced to a three-fold slowing in the degradation rate of the mutant (Fig. 3B). In contrast, the kinetics of disposal of immature WT CFTR were minimally altered by PIAS4; that is, a similar effect on ER associated degradation (ERAD) did not appear to account for increased expression of WT CFTR band B. The impact of PIAS4 to stabilize WT band C could obscure an action of the SUMO E3 on WT band B, since the ER is the source of folding-competent, mature CFTR at the PM.

Both *in vitro* and *in vivo* studies demonstrated that PIAS4 preferentially stimulated the SUMO-1 modification of F508del CFTR, while decreasing its conjugation to SUMO-2/3 (Fig. 5A&B). As SUMO-2/3 poly-chains are recognized by the ubiquitin E3, RNF4, we found also that PIAS4 reduced the ubiquitylation of F508del by 50% and eliminated the RNF4 induced reduction in the expression level of F508del CFTR.

PIAS4’s ability to promote CFTR biogenesis was striking; nevertheless, is PIAS4 a physiological modulator of CFTR biogenesis? Prior studies have primarily implicated PIAS proteins in their abilities to regulate nuclear events, usually based on their modulation of gene transcription (Rytinki *et al.,* 2009). qPCR did not detect significant PIAS4-induced changes in CFTR mRNA levels, nevertheless, it is possible that changes in the expression of other proteins contribute to the increases in immature CFTR that we observe for both WT and mutant CFTR proteins in response to PIAS4 expression. Other findings are consistent with a direct effect of the SUMO E3. First, when purified PIAS4 protein was added to *in vitro* SUMOylation reactions that included either SUMO-1 or SUMO-2, the modification of F508del NBD1 by SUMO-1 was nearly doubled, while its conjugation to SUMO-2 was significantly reduced (Fig. 5A). These results were reproduced *in vivo* when full-length CFTR, and its SUMO-1 modification were increased by PIAS4 over-expression in CFBE-F508del cells (Fig. 5B). In addition, the knockdown of PIAS4 as well as expression of the F508del-4KR mutant CFTR having mutations in four consensus SUMOylation sites provided evidence that this pathway, which augments ER-based CFTR biogenesis, regulates CFTR directly. The changes in CFTR expression evoked by PIAS4 were also mimicked by the expression of activated SUMO-1 or SUMO-2, conditions under which the effects of the SUMO E3 were reproduced in the absence of exogenous PIAS4 expression. While we cannot rule out the possibility that epigenetic changes contribute to the impact of PIAS4 on CFTR biogenesis, the ability to reproduce the impact of PIAS4 *in vivo,* with activated SUMO paralogs and with purified proteins *in vitro* make this explanation less likely.

### The SUMO Paralog Switch

Our findings reflect the presence of opposing pathways that determine the fate of CFTR, biogenesis vs. degradation, mediated by its post-translational, and perhaps co-translational, conjugation with different SUMO paralogs, SUMO-1 or SUMO-2/3. A similar divergence in functional outcome based on SUMO paralog exchange at target protein conjugation sites has been termed a ‘paralog switch’ (Fasci *et al.,* 2015). This behavior has been documented for the promyelocytic leukemia protein, PML, and its chemotherapeutic response to arsenic trioxide treatment. Arsenic reorganizes PML into nuclear bodies, from which it is targeted for degradation by the STUbL, RNF4. Arsenic trioxide induces the exchange of SUMO-2, conjugated with Lys^65^ of PML, for SUMO-1, which also drives SUMO-2 conjugation and SUMO chain extension at the downstream site, Lys^160^. The resulting SUMO-2 poly-chains are accessed by RNF4, leading to PML ubiquitylation and its proteasomal degradation. Arsenic treatment is thought to lead to PML oxidation, altering its conformation, which is thought to determine the timing of the signal associated with these site-specific paralog modifications.

Although not involving a functional paralog switch, a protease controlled paralog specific modification has been observed for the GTPase activating protein for Ran, RanGAP1, the first recognized SUMO substrate in mammalian cells. This GAP protein is conjugated to SUMO-1 and SUMO-2 with identical efficiency *in vitro. Yet in vivo,* RanGAP1 modified with SUMO-1 exhibits a higher affinity for its SUMO E3 ligase, RanBP2/Nup358, and interaction of the SUMO E2, Ubc9, with this complex provides protection from SENP protease activity which could otherwise remove SUMO-1. Thus, RanGAP1-SUMO-2 remains exposed and is efficiently cleaved, yielding RanGAP1-SUMO-1, the predominant form maintained *in vivo* (Zhu *et al.,* 2009; Gareau *et al.,* 2012).

In a similar protease controlled mechanism, paralog selectivity might be generated for WT and mutant CFTR. Cells express an excess of SUMO-2/3, which is maintained in a free pool; in contrast, the majority of cellular SUMO-1 is conjugated to substrates (Flotho and Melchior, 2013). If this is true also for epithelial cells expressing CFTR, these findings would suggest that both WT and F508del CFTR might be conjugated to SUMO-2/3, a default process that would promote degradation unless a stable CFTR conformation is achieved. Proteases cleaving off SUMO-2/3 might effectively reverse this SUMO modification of WT CFTR, permitting a shift toward SUMO-1 and promoting WT CFTR maturation. Folding deficiencies associated with mutant CFTR might obviate its access to SUMO protease and prevent SUMO-2/3 cleavage, directing F508del CFTR towards degradation.

The order of paralog addition to CFTR could also be opposite to that suggested above. The modification of CFTR by SUMO-1 may occur early, as a means of stabilizing nascent CFTR during its protracted domain folding/assembly process, which may take 30 min or more (Du *et al.*, 2005). Several lines of evidence favor the concept that failure of the F508del mutant to achieve appropriate domain-domain interfaces is the limiting factor in mutant protein progression (Sharma *et al.*, 2004; Du *et al.*, 2005; Lewis *et al.*, 2005; Thibodeau *et al.*, 2005; Cui *et al.*, 2007; Rabeh *et al.*, 2012). SUMO modification is recognized to increase protein solubility and reduce aggregation (Butt *et al.*, 2005; Esposito and Chatterjee, 2006). Indeed, commercial kits (e.g. www.lifesensors.com) feature SUMO fusions for generating difficult-to-express proteins (Lee *et al.*, 2008), and a SUMO fusion construct has been used to optimize the production of CFTR NBD1 and NBD2 for *in vitro* studies (Qu *et al.*, 1997). Likewise, our findings suggest that Nature might utilize this strategy for stabilizing difficult-to-fold proteins. F508del is an intractable protein that stalls at one or more conformational transitions that must be overcome energetically and kinetically to achieve the native fold (Du and Lukacs, 2009). Thus, SUMO-1 could assist in stabilizing immature CFTR, but if this process ultimately fails, a switch of SUMO-1 to SUMO-2/3, perhaps protease mediated, would expose the mutant to recognition by pathways that culminate in its disposal. The delineation of mechanism(s) that mediate this paralog switch should clarify this decision point for manipulations that could therapeutically modulate CFTR fate.

## MATERIALS AND METHODS

### Antibodies and Reagents

Monoclonal antibodies targeted CFTR NBD1, NBD2 and R-domain (#660, #596 and #217, respectively; Cystic Fibrosis Foundation Therapeutics, Bethesda, MD); other monoclonal antibodies were to the HA- or Flag-tag (Sigma-Aldrich, St. Louis, MO). Polyclonal antibodies to ubiquitin, Hsp70, SUMO-1 and SUMO-2/3 were from Enzo life Sciences (Ann Arbor, MI), antibodies to actin, PIAS4 and the myc-tag were obtained from Sigma-Aldrich (St. Lois, MO). Horseradish peroxidase-conjugated secondary antibodies, anti-mouse and anti-rabbit were obtained from Amersham-Pharmacia Biotech (GE Healthcare Bio-Sciences, Piscataway, NJ) and *N*-Ethylmaleimide (NEM) was purchased from Thermo Scientific (Rockford, IL), complete protease inhibitor cocktail tablets were obtained from Roche Diagnostics Corporation (Indianapolis, IN), and other chemicals were obtained from Sigma-Aldrich (St. Louis, MO) at highest grade available.

### Plasmid constructs, Adenoviral vector and Site-directed mutagenesis

The pCMV-Flag-hPIAS4 and pCMV-Flag-hPIASx alpha plasmids were obtained from Addgene (Cambridge, MA). The Flag-RNF4 plasmid was a generous gift of Dr. Ronald T. Hay (University of Dundee, Dundee, UK). The pCMV-Flag-hPIAS4-CA (C342/347A) and pCMV-Flag-hPIAS4-2KR (K35/128R) mutants were generated using the following primers: 5’-CCGGGCAGAGACCGCCGCCCACCTGCAG-3’ (C342A, forward), 5’-CTGCAGGTGGGCGGCGGTCTCTGCCCGG-3’ (C342A, reverse), 5’-CGCCCACCTGCAGGCCTTCGACGCCGTC-3’ (C347A, forward), 5’-GACGGCGTCGAAGGCCTGCAGGTGGGCG-3’ (C347A, reverse).

The pcDNA3.1-F508del CFTR-4KR (K377/447/1199/1468R) mutant was generated using the primers: 5’-TGGAGCAATAAACAAAATACAGGATTTCTTACAAAGGCAAGAATATAAGACATT-3’ (K377R, forward), 5’-AATGTCTTATATTCTTGCCTTTGTAAGAAATCCTGTATTTTGTTTATTGCTCCA-3’ (K377R, reverse), 5’-CCTGTCCTGAAAGATATTAATTTCAGGATAGAAAGAGGACAGTTG-3’ (K447R, forward), 5’-CAACTGTCCTCTTTCTATCCTGAAATTAATATCTTTCAGGACAGG-3’ (K447R, reverse), 5’-TTGAGAATTCACACGTGAGGAAAGATGACATCTGGCC-3’ (K1199R, forward), 5’-GGCCAGATGTCATCTTTCCTCACGTGTGAATTCTCAA-3’ (K1199R, reverse), 5’-AGCCCCAGATTGCTGCTCTGAGAGAGGAGACAG-3’ (K1468R, forward), 5’-CTGTCTCCTCTCTCAGAGCAGCAATCTGGGGCT-3’ (K1468R, reverse).

Mutants were obtained using the QuickChange lightning Site-Directed Mutagensis Kit (Agilent Technologies, Santa Clara, CA). All plasmid constructs were sequence verified.

For adenovirus production, PIAS4 was amplified by RT-PCR from a CFBE cDNA library. The amplification fragment was cloned into the pAD/DEST vector (Invitrogen, Carlsbad, CA), the clones were confirmed by sequencing, and DsiRNA targeting PIAS4 were synthesized from literature experience (Morris *et al.*, 2009) and obtained from Integrated DNA Technologies (Coralville, Iowa). The DNA cloned into the pAD/CMV/DEST vector was then confirmed by sequencing (Genewiz, New Jersey, NY) and 293A cells were transfected with pAd/CMV/V5-DEST-GFP-PIAS4 or its -DsiRNA using Lipofectamine 2000 (Invitrogen). High titer adenovirus stocks were generated from 293A cell crude homogenates using the Vivapure AdenoPACK 100 kit following the manufacturer’s instructions.

### Cell culture and protein expression

CFBE41o- airway cells were cultured in Minimal Essential Medium (Invitrogen, Carlsbad, CA) and supplemented with 10% Fetal Bovine Serum (Hyclone, Logan, UT), and 2 mM L-glutamine, 50 U/ml penicillin and 50 µg/ml streptomycin. For CFBE WT and F508del stable cell lines, 0.5 µg/ml and 2 µg/ml Hygromycin B (InvivoGen, San Diego, CA) were added respectively as selection agents. All cells were maintained in a humidified chamber with 5% CO_2_ at 37°C

For protein over-expression, CFBE-WT, -F508del stable cell lines (Bebok *et al.*, 2005) or parental CFBE41o-cells grown in 6-well plates were transiently transfected with the indicated expression plasmids using Lipofectamine 2000 (Invitrogen) as indicated. After 48 h, cells were rinsed with phosphate-buffered saline and lysed in RIPA buffer (50 mM Tris-HCl, pH 8.0, 150 mM NaCl, 1.0% Triton X-100, 0.5% sodium deoxycholate, 0.1% SDS) with protease inhibitors. Samples were incubated for 15 min in RIPA buffer followed by several short bursts of sonication with a tip sonicator, and then centrifuged at 16,000 × g for 10 min at 4°C. Cell lysates were used for immunoblot analysis.

### Immunoblotting and co-immunoprecipitation assays

Equal amounts of proteins from CFBE-WT, -F508del or parental cell lysates were resolved by SDS–PAGE and transferred to PVDF membranes (PerkinElmer, Boston, MA). Unbound sites were blocked for 1 h at room temperature with 5% non-fat milk powder in TBST (20 mM Tris, 150 mM NaCl, 0.01% Tween 20, pH 8.0). The blots were incubated with primary antibodies (anti-CFTR #217, 1:5,000; anti-actin, 1:4,000; anti-Hsp70, 1:2,000; anti-Flag, 1:10,000; anti-PIAS4, 1:1,000) at room temperature for 1 h. The blots were then washed three times for 10 min each with TBST and incubated for 1 h with horseradish peroxidase–conjugated secondary antibodies in TBST containing 5% non-fat milk, followed by three TBST washes. The reactive bands were visualized with enhanced chemiluminescence (PerkinElmer Life Sciences, Wellesley, MA) and exposed to X-ray film, or for some experiments, imaged and quantified on a Li-Cor Odyssey.

For co-immnunoprecipitation assays, pre-cleared cell lysates (~ 1mg of protein) were mixed with respective primary antibodies for 2 h at 4°C in lysis buffer (20 mM Tris, 150 mM NaCl, 10% glycerol, 1% Triton X-100, 2 mM EDTA, pH 8.0, containing protease inhibitors). For the detection of *in vivo* SUMOylation, 20 mM N-ethylmaleimide (NEM) was added to the lysis buffer. Fifty microliters of washed protein A- or G-agarose beads was added to each sample and incubated 4 h at 4°C with gentle rotation. Immunocomplexes were washed three times with lysis buffer and precipitated by centrifugation at 12,000 × g for 10 s. Then the immunocomplexes were resuspended in SDS sample buffer and subjected to immunoblotting.

### *in vitro* SUMOylation assays

The *in vitro* assay was performed using reagents purchased from Enzo Life Sciences (Ann Arbor, MI) together with F508del human NBD1 protein that contained a single solubilizing mutation, F494N, as previously described (Rabeh *et al.*, 2012; Gong *et al.*, 2016). In brief, 15 ng of purified NBD1 was incubated in SUMOylation buffer with a reaction mixture containing recombinant E1 (0.4 μM), Ubc9 (4 μM), a SUMO paralog (3 μM), and Mg-ATP (2 mM), with or without purified recombinant human PIAS4 protein (15 ng), for 1 h at 27°C. After the reaction was terminated with SDS sample buffer containing 2-mercaptoethanol, reaction products were fractionated on 12% SDS–PAGE, and the gel shift resulting from SUMO modification was detected by immunoblotting using anti-NBD1 (#660).

### Cycloheximide (CHX) Chase

CFBE-WT, -F508del or parental cells were plated and transiently transfected as described above. After 48 h, cells were incubated with 100 μg/ml CHX for the indicated times (Fig. 3), at which cells were harvested, lysed with RIPA buffer containing protease inhibitors and subjected to immunoblot to determine the time-course of CFTR decay following the inhibition of protein synthesis.

### *in vivo* Ubiquitylation assays

CFBE-F508del cells were plated and transiently transfected as described above. After 48 h, the cells were treated with 50 μM MG132 for 6 h, and then lysed in lysis buffer containing protease inhibitors. Pre-cleared lysates were mixed with CFTR antibodies (#217 and #596) for 2 h at 4°C in lysis buffer. 50 µl of washed protein G agarose beads was added to each sample and incubated 4h at 4°C with gentle rotation. Immunocomplexes were analyzed by immunoblotting with anti-Ubiquitin antibody.

### Total RNA extraction and Real-Time PCR

CFBE-F508del cells were transfected with empty vector, pCMV-Flag-hPIAS4 or pCMV-Flagh-PIASx alpha plasmids. After 48 h, total RNA from transfected cells was purified using the RNeasy Kit (Qiagen, Valencia, CA) according to the manufacturer’s instructions; 1 μg of the total RNA was converted to cDNA using the iScript cDNA Synthesis Kit (Bio-Rad Laboratories, Hercules, CA). Equal amounts of each single-chain cDNA were subjected to qPCR using Bio Rad iQ SYBR Supermix. The qPCR reaction was completed using the CFX96 Touch Real-Time PCR Detection System (Bio-Rad) for 3 min at 95°C followed by 39 cycles of 10 sec at 95°C and 30 sec at 55°C. The primer sequences used were: CFTR: 5’-CACAGCAACTCAAACAACTGG-3’ (forward), 5’-TGTAACAAGATGAGTGAAAATTGGA-3’ (reverse); actin: 5’-ATTGGCAATGAGCGGTTC-3’ (forward), 5’-CGTGGATGCCACAGGACT-3’ (reverse).

The Ct values were determined for each reaction (in triplicate) using Bio-Rad CFX Manager Software, and quantification was completed using the ΔΔCt method.

### Expression of functional cell surface protein

CFBE-WT or -F508del cells were subjected to cell surface biotinylation as described (Silvis *et al.*, 2009). Briefly, control cells or those transduced with AdPIAS4 were treated with EZ-Link Sulfo-NHS-LC-LC-Biotin (Pierce, Rockford, IL). The experiments were performed on ice with all solutions at 4°C. To biotinylate apical proteins, cells were rinsed once with PBS+CM (lacking CaCl_2_ + MgCl_2_) and then washed with PBS+CM containing 10% FBS. The apical surface was biotinylated with 1 mg/ml EZ-Link Sulfo-NHS-SS-Biotin in borate buffer (85 mM NaCl, 4 mM KCl, and 15 mM Na2B4O7, pH 9) for 30 min with gentle agitation. Excess biotin was removed using two 10-min washes in PBS+CM with 10% FBS, followed by two washes in PBS+CM.

Cell surface CFTR was quantified by lysing cells in 700 μl biotinylation lysis buffer (BLB: 0.4% deoxycholate, 1% NP40, 50 mM EGTA, 10 mM Tris-Cl, pH 7.4, and Complete EDTA-free Protease Inhibitor Cocktail, Roche Diagnostics). After protein levels were determined, normalized samples were incubated overnight with 150 μl UltraLink Immobilized NeutrAvidin Protein Plus (Pierce). Precipitated proteins were washed three times with BLB, solubilized with Laemmli sample buffer, resolved by SDS-PAGE and blotted for CFTR.

A clone of CFBE cells stably expressing a FAP-F508del CFTR construct was generated as previously described for a 293A stable cell line (Larsen *et al.*, 2016). These FAP-CFBE-F508del expressing cells were plated in a 96-well plate in complete DMEM containing 8 µg/mL polybrene with or without addition of recombinant adenovirus encoding PIAS4. Two days post plating, the growth media was exchanged for complete DMEM containing 2 µM VX-809 and cells were transferred to a tissue culture incubator set at 27°C. The following day, FAP-F08del CFTR surface expression was detected quantitatively by automated microscopy analysis as described previously (Larsen *et al.,* 2016).

Whole-cell currents were recorded from transfected CFBE41o- cells as previously described for HEK293 cells (Bertrand *et al.,* 2009). Briefly, the electronics consisted of a 200B Axopatch amplifier controlled by Clampex 8.1 software through a Digidata 1322A acquisition board (Axon Instruments). Solutions were maintained at 37°C with a flow rate of 2.0 ml/min. The imaging chamber was mounted on the stage of a Nikon Diaphot microscope equipped with standard illumination and a xenon lamp with GFP filter cube (Ex 485/Em 550) permitting identification of GFP-expressing cells. Seal resistances exceeded 8 GΩ and pipette capacitance was compensated; all experiments used a standard voltage-clamp protocol with a holding potential of −40 mV. NMDG-Cl was used for both the bath and pipette solutions to isolate chloride currents. The bath solution was (in mM): 140 NMDG-Cl, 10 HEPES, 1 MgCl_2_, 1.5 CaCl_2_, 5 glucose, pH 7.3, and the pipette solution was (in mM): 140 NMDG-Cl, 10 HEPES, 1 MgCl_2_, 5 glucose, 1 EGTA, pH 7.2 and also contained 1 mM Mg-ATP and 100 μM GTP. The osmolarity of the bath and pipette solutions were measured and NMDG-glutamate was added to the bath solution to generate a 25 mOsm gradient that was needed to obviate cell swelling (Worrell *et al.,* 1989). Pipettes using thin wall borosilicate glass were pulled to tip diameters of 1–2 μm (access resistance < 4 MΩ).

### Statistical Analysis

A P-value of <0.05 was considered statistically significant. The normality of data distribution was validated by the Kolmogorov-Smirnov normality test and visual inspection of quantile-quantile (QQ) plots. Statistical analysis was performed using the paired t–test with the means of at least three independent experiments if the data were normally distributed. If this condition could not be confirmed, the Wilcoxon signed-rank test was used. We used SAS (SAS Institute Inc., Cary, NC, USA) for statistical analyses.

## ACKNOWLEDGMENTS

This work was supported by grants from the National Institutes of Health (DK68196 and DK72506) and the Cystic Fibrosis Foundation (FRIZZE05XX0). Thanks to Dr. Marcel Bruchez of Carnegie Mellon University for supplying MG-B-Tau for the FAP experiments and to Dr. Gergely Lukacs, McGill University, Montreal, CA for supplying F508del NBD1 with a single solubilizing mutation.

